# ProtView: A versatile tool for *in silico* protease evaluation and selection in a proteomic and proteogenomic context

**DOI:** 10.1101/2021.09.02.458698

**Authors:** Sophia S. Puliasis, Dominika Lewandowska, Piers Hemsley, Runxuan Zhang

## Abstract

Tools have been created to generate *in silico* proteome digests with different protease enzymes and provide useful information for selecting optimal digest schemes for specific needs. This can save on time and resources and generate insights on the observable proteome. However, there remains a need for a tool that evaluates digest schemes beyond protein and amino acid coverages in the proteomic domain. Here, we present ProtView, a versatile in-silico protease/protease combination and digest evaluation workflow that maps *in silico* digested peptides to both protein and genome references, so that the potential observable sections of the proteome, transcriptome and genome can be identified. This supports the identification and quantification of the proteomic evidence of transcriptional, co-transcriptional, post-transcriptional and translational regulations. Benchmarking against biological data comparing multiple proteases shows that ProtView can correctly estimate the relative performances among the digest schemes. ProtView provides this information in a way that is easy to interpret, allowing for digest schemes to be evaluated before carrying out an analysis, in a broader context to optimize proteomic and proteogenomic experiments. ProtView is available at https://github.com/SSPuliasis/ProtView.

## INTRODUCTION

Bottom-up proteomics involves using proteases to digest protein mixtures into peptides, which are then analysed by mass spectrometry (MS), allowing the peptides, and therefore the originating protein, to be identified (Aebersold and Mann, 2003). Shotgun proteomics refers to the use of bottom-up methods to identify proteins in complex mixtures. The shotgun proteomics workflow typically begins with the protein sample being denatured, reduced, alkylated, and digested by one or more proteases into peptides, which are then separated by liquid chromatography and identified by tandem mass spectrometry (MS/MS) and database searching (Wu et al., 2002). Database searching determines whether a peptide sequence in a database gives a significant match to each MS/MS spectrum and the degree of matching is assigned a score (Cottrell, 2011). It is paramount that every step is carried out effectively to maximise peptide identification and quantification, eventually maximising protein coverage and quantification. When proteins are digested in the first stage, it is the resulting peptides that are carried into the subsequent analysis. Therefore, a peptide not generated in the digest cannot be identified in the subsequent analysis.

Trypsin is usually the protease of choice because it is highly specific, cleaving C-terminal to Lysine and Arginine residues (Keil et al., 1992), stable under a wide range of experimental conditions, and generates peptides in the preferred low charge and 7-35 amino acid length range for detection by MS machinery (Swaney et al., 2010), although it is not always the most suitable choice. For example, lysine and arginine are less frequent in membrane spanning protein regions (Kyte and Doolittle, 1982), resulting in fewer detectable peptides per unit length, thus limiting identification and quantification using MS methods when membrane spanning regions are digested with trypsin. Lysine and arginine are also enriched at exon-ending and junction residues due to their codons (Wang et al., 2018), resulting in trypsin cleavage at splice junctions. This impedes the detection of junction-spanning peptides necessary to identify splice isoforms arising from alternative mRNA splicing. Furthermore, due to enzyme specificity, the perpetual use of any highly specific protease will continuously generate the same sub-sets of peptides. This eventually leads to a ‘tunnel vision’ display of the proteome in databases and repositories, with regions or even whole proteins that do not produce MS/MS suitable peptides with the protease in question being unidentified and remaining uncharacterised (Tsiatsiani and Heck, 2015).

Digests with different proteases, either to replace or to complement trypsin, have emerged as a way of mitigating the above issues and have proven to be useful in the study of membrane proteins (Fischer and Poetsch, 2006), splice junctions (Wang et al., 2018), N-termini not accessible by trypsin (Soh et al., 2020), and the study of post-translational modifications (PTMs) (Tran et al., 2016). Explorations into alternative enzymes support the argument that there could be an ideal, non-trypsin centric, digestion scheme for every biological question and type of analysis (Tsiatsiani and Heck, 2015).

Expanding upon this idea, multiple protease digestion strategies can also be brought to bear on the issue of uncharacterised proteins. Combining multiple enzymes can be done in parallel, where peptide information from single protease digests is combined during post MS/MS analysis, or concurrently, where multiple proteases are added to the same sample in vitro before MS/MS analysis is carried out. It has been reported that using enzymes in parallel results in a significant increase in sequence coverage compared to single digests of the *Saccharomyces cerevisiae* proteome (Swaney et al, 2010) *Cannabis sativa* buds (Vincent et al., 2019a, Vincent et al., 2019b), human cervical cancer cells (Guo et al., 2014), and human recombinant protein (Choudhary et al, 2003). On the other hand, concurrent digests were reported to increase the number of identified proteins when Trypsin-Asp-N was used on *Schizosaccharomyces pombe* whole cell lysates when compared to trypsin alone (Dau et al., 2020), and be more efficient at yielding fully cleaved peptides and reducing the abundance of missed cleavage in peptides when Trypsin-Lys-C were used on *S. cerevisiae* (Glatter et al., 2012).

The aforementioned studies can make digest scheme recommendations for specific species and types of experiment after carrying out comparisons between digest schemes *in*-*vitro*. The scope of proteomic analyses is very broad and knowing which digest scheme is better suited to an analysis beforehand can save on costs, time and resources. Programs such as PeptideCutter (Gasteiger et al., 2003) and Rapid Peptides Generator (RPG) (Maillet, 2019) can digest protein sequences with different enzymes *in-silico* to give peptides that will theoretically be generated by a digest. ProteaseGuru (Miller at al., 2021) and Proteogest (Cagney et al., 2003) go a step further and provide interpretations of their digest results to aid in protease selection. Proteogest is a Perl application that allows the user to select a combination of provided or custom modifications, to assess the effects of PTMs on the outcome. ProteaseGuru is a versatile and accessible tool that provides detailed peptide information and includes database PTM annotations and data visualization in the outputs. Nonetheless, there remains a need for such a tool that can also provide information in a wider context, that includes transcriptomic and genomic coverages and regions, e.g. to aid the study of alternative splicing and identify peptides that are unique to individual transcript isoforms, thus allowing the identification and quantification of the effects of transcriptional and translational regulations.

This work introduces ProtView, a method that integrates *in-silico* digestions by Rapid Peptides Generator (RPG) and provides the set of possible peptides that can be identified by each protease, or protease combination, and the variable information that they present, such as peptide length distributions, protein sequence coverage, and amino acid coverage. It also maps the digested peptides back to the genome using coding sequence (CDS) information from the annotations, which can provide detailed locations and information of the digested proteome in transcriptomic and genomic context, enabling analyses such as the identification of splice junction covering peptides, and isoform-unique peptides. It is the first tool that allows for the use of *in silico* proteomic evidence to investigate transcriptional and translational regulations to study the proteomic impact of alternative splicing, polyadenylations, alternative transcriptional starting locations, as well as alternative translational site regulations. We have demonstrated the utility of ProtView, with an analysis on the *Arabidopsis thaliana* proteome. We also compared ProtView predictions with *in-vitro* protease experiments and published proteomic data.

## METHODS

ProtView is a novel computation tool that presents all the *in silico* digested peptides by each protease, or protease combination, and their detailed information on the proteome, transcriptome, and genome, such as peptide length distributions, protein sequence coverage, specific residue coverage, peptides that cover splice junctions, junction coverage, the number of isoform-unique peptides, and genomic coordinates of peptides (Figure 1). All programming was done in Python 3.8 under the GPL v3 license. Details of the program and instructions can be found at (https://github.com/SSPuliasis/ProtView).

**Figure 1.**
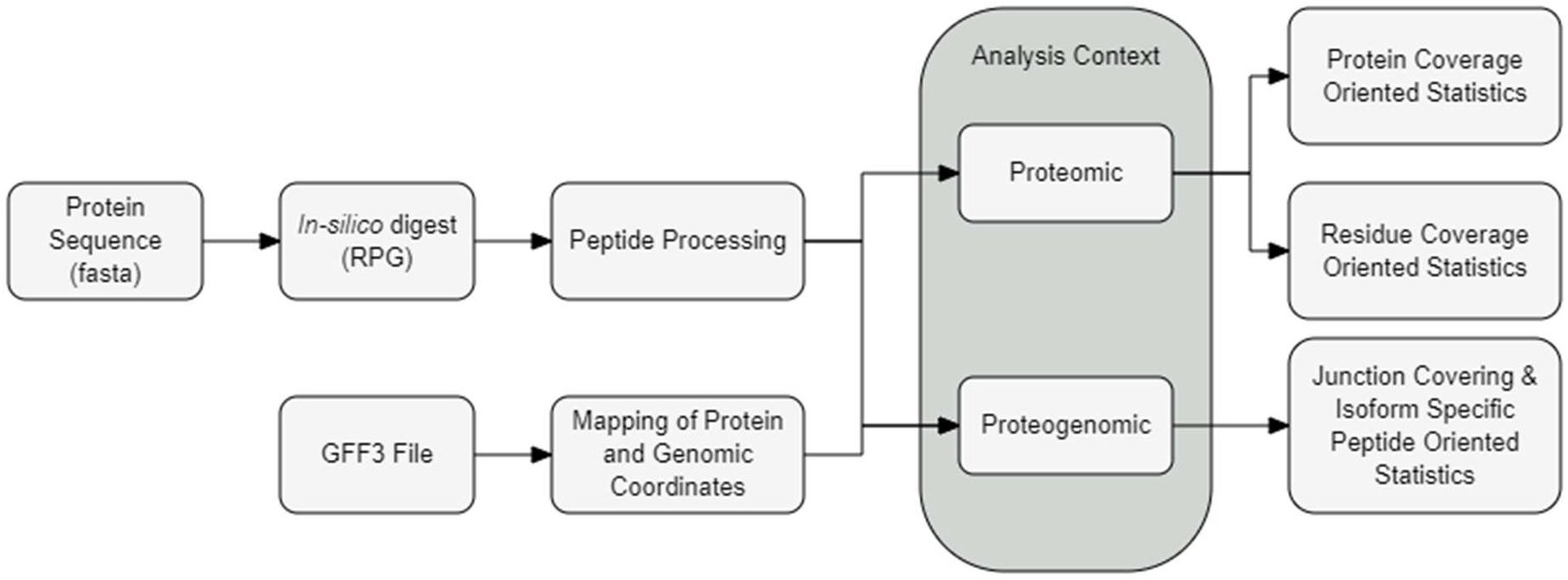
Outline of the ProtView workflow

## PROTVIEW WORKFLOW

### Peptide Processing

Rapid Peptides Generator (RPG) (Maillet, 2019) is incorporated into ProtView because it can process the whole protein database in one go, allows for user-defined proteases, and can generate more information than PeptideCutter (Gasteiger *et al*., 2005), such as isoelectric point of each peptide. RPG has the option to carry out single or concurrent digests, where a sequence is simultaneously cleaved by multiple enzymes. Based on the output of RPG, ProtView creates parallel enzyme digests by combining peptides from individual single digests.

Duplicates of peptides that are generated by more than one of the single digests (e.g., by proteases with similar cleavage specificities) are removed from the parallel digest, resulting in a non-redundant parallel digest output. The enzyme name in the parallel digest output is the enzymes used separated by ‘/’, whereas the enzyme name for concurrent digests in RPG is the enzymes used separated by ‘-’. Peptides can then be filtered by amino acid length and/or by content of a specific residue. Filtering for a length of 7-35 amino acids is recommended as the detectable range, although this optional filtering can be set by the user to match the instrumentation or criteria for the experiment. Below shows an example of a peptide generated by RPG and processed by ProtView (Table 1).

**Table 1.** Example of a digested peptide after processing digest results. The table contains information on the enzyme used in the theoretical digest, relative peptide start and cleavage position (end) on the protein sequence, size in amino acids, molecular weight, isoelectric point, peptide sequence, and the protein that the peptide originates from

| enzyme | Peptide start | Peptide End | Peptide size | Mol weight | Isoelectric point | Sequence | Parent |
| --- | --- | --- | --- | --- | --- | --- | --- |
| Trypsin | 1 | 8 | 8 | 991.1297 | 7.77 | MFSNIDHK | AT1G66600.1 |

### Proteomic Summary Statistics

Protein sequence coverage is the percentage of the original protein sequence digested that is covered by filtered peptides and is calculated as the ratio between the sum of filtered peptide lengths and the total lengths of protein sequences in the FASTA file. Residue coverage is the percentage of an amino acid in the original FASTA sequence file that is covered by the filtered peptides. The total number of peptides generated by each enzyme both before and remaining after filtering are also presented in the table, alongside their mean and median lengths. Table 2 is a summary table generated by ProtView for a single gene as an example, expandable to multiple genes or a proteome.

**Table 2.** Summary statistics of a digested protein after processing digest results, with Threonine(T) coverage

| enzyme | Total peptides | Mean length | Median length | Filtered peptides | Sequence coverage (%) | T coverage (%) |
| --- | --- | --- | --- | --- | --- | --- |
| Asp-N | 95 | 8.9 | 7 | 46 | 70.7 | 81.2 |
| Glu-C | 104 | 8.1 | 5 | 43 | 80.6 | 96.9 |

### Coding Sequence (CDS) preparation for downstream analyses

When the genomic coordinate for each protein is available in gene and transcript annotations as coding sequences (CDS) in gff3 format, ProtView assigns a unique ID to each CDS and the adjacent intron preceding it, calculates the length of each intron between CDS, and converts the genomic coordinates of CDS regions to the relative protein sequence coordinates. CDS and intron IDs consist of chromosome, start position, end position, and strand (e.g., Chr1_24848737_24848859_+). While converting CDS genomic coordinates to their relative proteomic coordinates, intron lengths between CDS are added up separately for each isoform, to give cumulative intron length of the introns preceding each CDS within an isoform. To obtain the relative protein coordinates for a CDS on the positive strand, the translation start position of the gene and cumulative intron length (after translation start position) are subtracted from the CDS genomic coordinate increased by one, then divided by 3 to take the triplet code into account. The final number is rounded up to the nearest integer as shown in the equation below. While for CDS on the negative strand, the same equation can be applied with adjustments shown below.

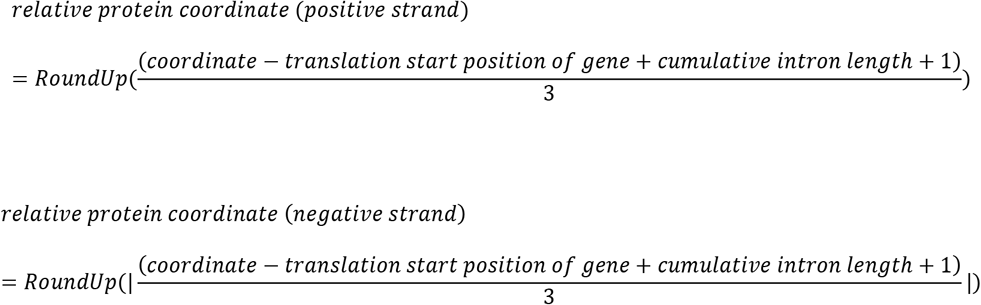

Output from this step is saved as two csv files, one for each DNA strand, and allows for the downstream conversion of relative proteomic peptide coordinates to genomic and the identification of peptides covering splice junctions. Table 3 shows an example of the output table for a protein on the positive strand.

**Table 3.** Format of CDSs after processing and extraction from GFF3 files

| type | start | end | strand | Parent | CDS ID | Intron ID | Intron length | Cumulative intron length | Protein start | Protein end |
| --- | --- | --- | --- | --- | --- | --- | --- | --- | --- | --- |
| CDS | 24848737 | 24848859 | + | AT1G66600.1 | Chr1_24848737_24848859_+ | Chr1_24848651_24848736_+ | 86 | 86 | 83 | 123 |

### Genomic coordinates of the peptides

Relative peptide coordinates from the digest output are converted to the outer bounds of their corresponding coordinates on the genome, depending on DNA strand (Figure 2). The digested peptides are compared to CDS on the protein coordinates first, and the information from CDS that overlap with the digested peptide is used to calculate the genomic coordinates of the peptide. The conversion is shown as the equations below:

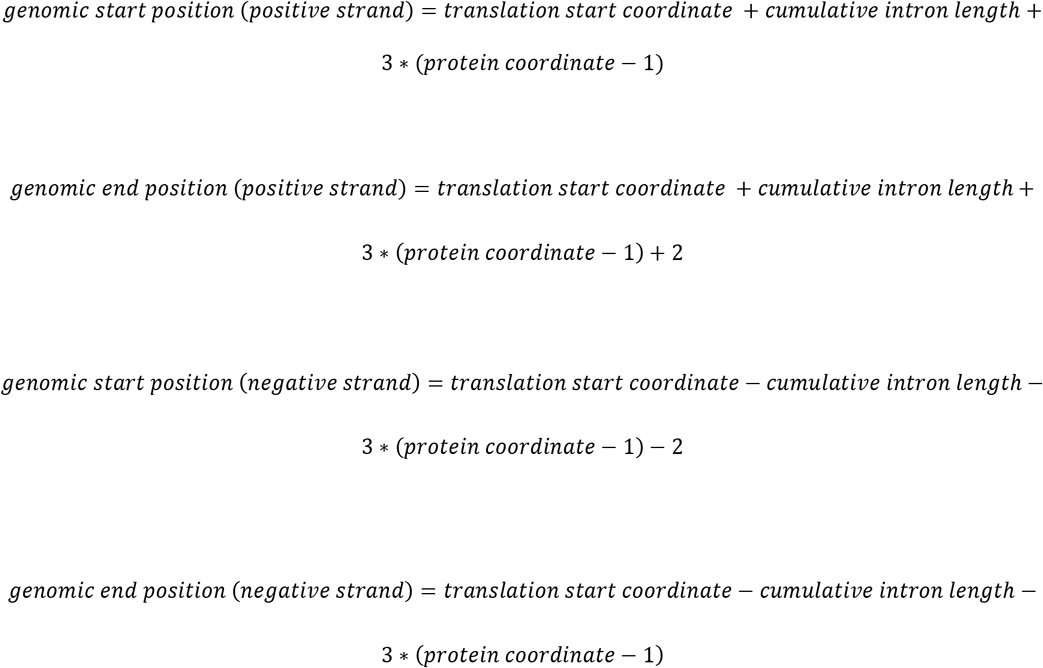

**Figure 2.**
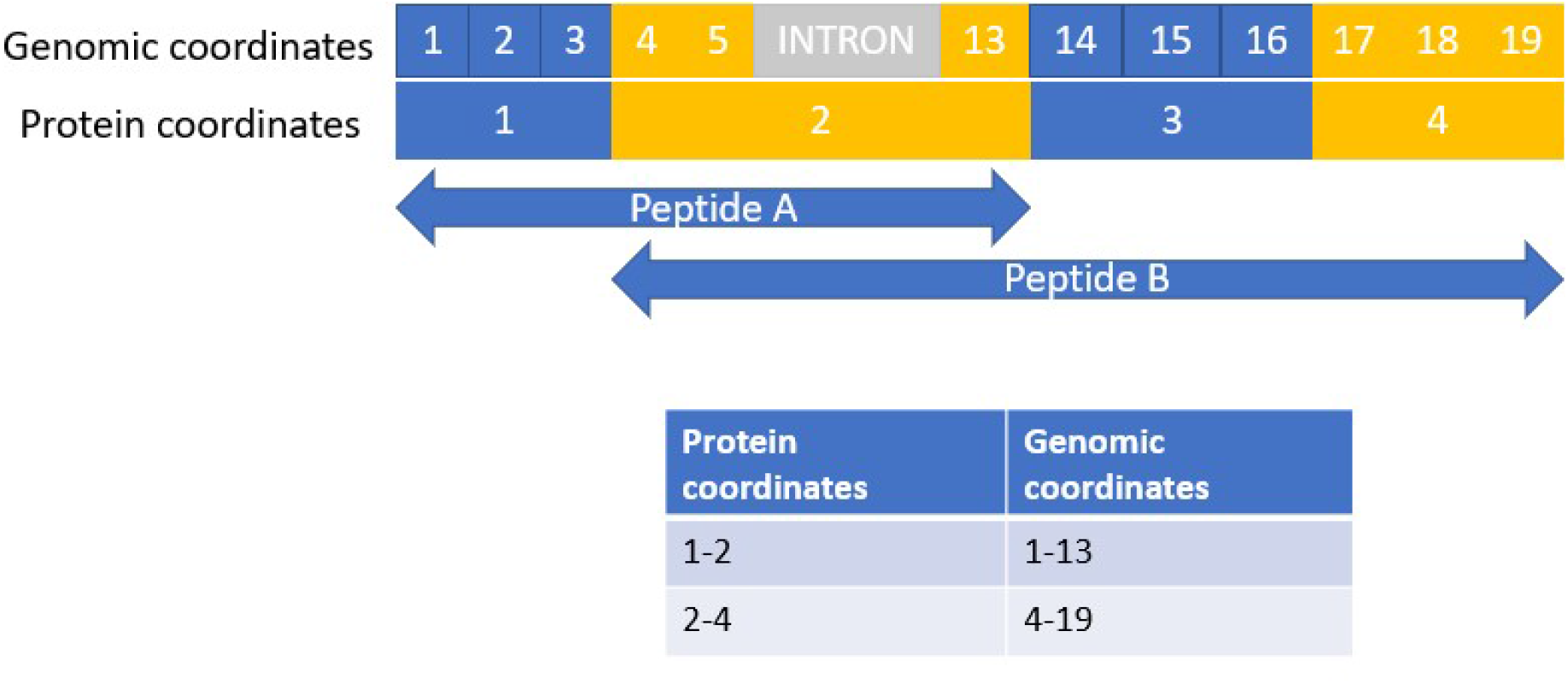
Illustration example of relative peptide coordinates and their corresponding coordinates on the genome. The top row of numbers represents nucleotides in the genome, with each triplet corresponding to one amino acid in the protein sequence. The row underneath represents the relative protein coordinates in amino acids (alternating blue and yellow).

In the example figure below (Figure 2), amino acids are colored in blue and yellow, and introns in grey. Nucleotide coordinates are numbered within amino acids, with three nucleotides corresponding to one amino acid. Peptides A and B each have the second amino acid as a coordinate, however the number given for the genomic coordinates differs between the peptides due to this amino acid representing the start of peptide B and the end of peptide A, and the script giving outer bounds of genomic positions.

The resulting data frame contains the parent isoform, both genomic and protein coordinates for each peptide, and the enzymes used to generate the peptide. Table 4 shows an example of two peptides (SPPVYRTTYLGQHTCKAFGVHD, DNTYGSEMINF), generated by digestion of AT1G66600.1 with Glu-C and Asp-N respectively, and their calculated genomic coordinates. Note that the peptides share the relative protein coordinate 169, but that their genomic coordinates differ after the conversion due to ProtView giving the outer bound of genomic coordinates, depending on whether a coordinate is at the start or the end of a peptide. Genome browser and visualization tools such as the R package Gviz (Hahne and Ivanek, 2016) can be used to visualize peptides mapped onto the genome, as shown in Figure 3.

**Table 4.** Output format of genomic coordinate conversion function of two exemplar digested peptides

| Peptide | isoform | Protein start coordinate | Genomic coordinate start | Protein end coordinate | Genomic coordinate end | enzyme |
| --- | --- | --- | --- | --- | --- | --- |
| SPPVYRTTYLGQHTCKAFGVHD | AT1G66600.1 | 148 | 24849051 | 169 | 24849116 | Glu-C |
| DNTYGSEMINF | AT1G66600.1 | 169 | 24849114 | 179 | 24849146 | Asp-N |

**Figure 3.**
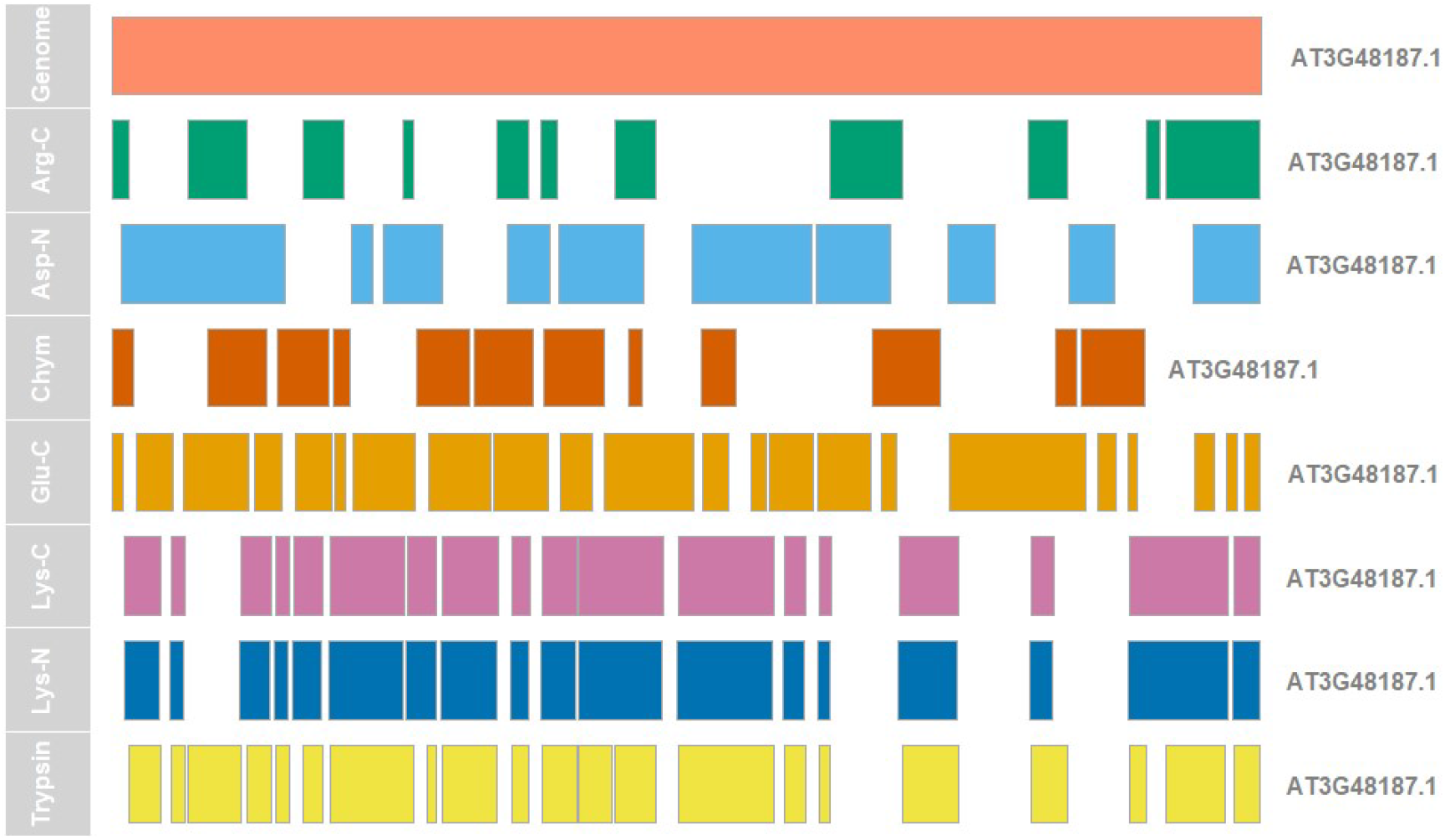
Gviz visualization of genome coverage for the mono-exonic gene AT3G48187.1 using peptides in the 7-35 aa range. The top row represents the genomic sequence. Regions of peptide sequence coverage generated by different proteases are shown in the subsequent rows.

### Junction covering peptide identification & statistics

Another function of ProtView is to identify peptides that span splice junctions. Relative protein splice junction coordinates are determined as the average of the CDS protein coordinates on each side (i.e. 20.5 if a upstream CDS ends at 20 and the downstream begins at 21). For each transcript/proteome isoform, each peptide is then checked against the junction coordinates and is considered as junction-covering if it has at least one amino acid encoded on either side of a splice junction. Positive outcomes are saved in the same format as the digest results, with an additional column for junction location. A summary table is generated from the junction-covering peptides, which includes the number of junction-covering peptides generated by each enzyme, the number of total and unique junctions that a digest scheme covers (to avoid double counting of splice junctions shared between transcripts), and junction coverage percentage, which is the percentage of the total junctions in the transcript isoforms being examined that are covered by digested peptides. Table 5 shows an example junction summary table.

**Table 5.**
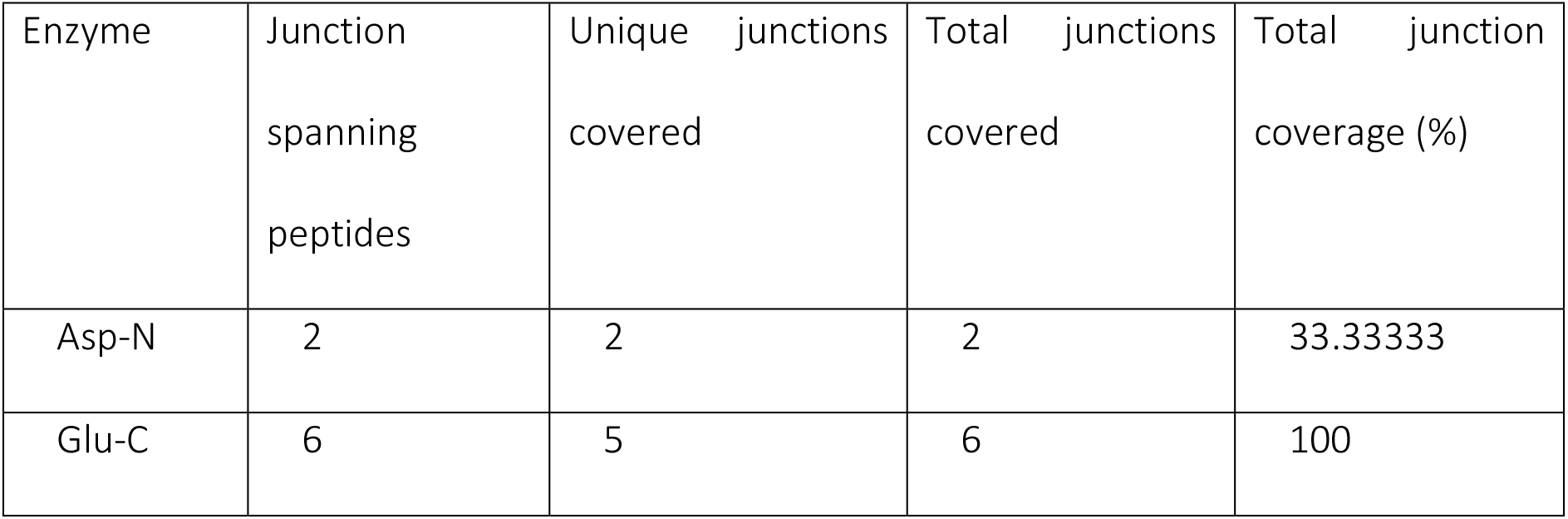
Junction summary statistics of digested proteins AT1G66600 and AT1G66610

### Isoform-unique peptides

Isoform-unique peptides can be used to identify a protein isoform with certainty and to discriminate between protein/transcript variants with different functions. The isoform-unique peptides come from exons that cover unique regions 1) due to alternative splicing events; 2) N or C terminus of the proteins resulting from different translation start or termination sites. The number of isoform-unique peptides is calculated by removing duplicate peptide sequences that can be found in more than one isoform from the filtered peptides, giving the number of peptides generated that can only be found in one isoform in each specific digest.

### Data acquisition & preparation

The Araport11 *A. thaliana* proteome (Cheng et al., 2017) is used in this manuscript as a dataset to illustrate the range of information that can be given by ProtView. The Arabidopsis protein sequence databases were downloaded from the TAIR database (https://www.arabidopsis.org/download/index-auto.jsp?dir=%2Fdownload_files%2FSequences%2FAraport11_blastsets), and corresponding GFF3 files from Araport11 (https://www.araport.org/data/araport11).

We have also used a number of additional published proteomics data that uses different enzymes for digestion to check the consistency with predictions with ProtView. The Human recombinant DNA derived tissue plasminogen activator protein sequence and *S. cerevisiae* proteome were downloaded from Uniprot (https://www.uniprot.org/uniprot/P00750, https://www.uniprot.org/proteomes/UP000002311) and digested using the same enzymes as the publications. All *in silico* digests were carried out using Rapid Peptides Generator (RPG) (Maillet, 2019) and filtered with ProtView for a recommended length of 7-35 amino acids.

## RESULTS AND DISCUSSION

### Using ProtView to analyse digests in a proteomic context

#### i. Selecting enzymes with the highest protein coverage

The purpose of the ProtView tool is to guide the choice of protease in *in vitro* experiments, by providing various statistics on peptides generated *in silico*. One of these measurements is the percentage of the original protein sequence that is covered by the digested and potentially identifiable peptides. While the coverage values given by ProtView are expected to be higher than those obtained in the laboratory, due to the theoretical analysis giving the upper limit of all possible peptides that could be identified, ProtView is useful in being able to show how digest schemes perform relative to one another.

The *A. thaliana* proteome was digested *in silico* by Rapid Peptides Generator (RPG) with Arg-C, Asp-N, Chymotrypsin (high specificity), Glu-C, Lys-C, Lys-N, Trypsin, and pairwise combinations of these proteases in concurrent (represented in this article by ‘-’) and parallel digests (represented in this article by ‘/’). The resulting peptide data will be used as an example to demonstrate the utility of ProtView. These proteases were chosen because they are highly specific, common alternative proteases, and often paired with trypsin in the literature. Due to the number of possible combinations, this example includes Trypsin in concurrent and parallel combination with Asp-N, Chymotrypsin, and Lys-C. Results for the remaining combinations can be found in the supplementary material. Summary statistics for these digests were calculated using ProtView and can be seen in Table 6, which includes the total number of peptides before and after filtering by length (7-35aa length used here), median and mean lengths of the peptides generated, the number of isoform-unique peptides, and sequence coverage %.

**Table 6.** Summary statistics of in silico digests carried out on A. thaliana sorted from highest to lowest protein sequence coverage %

| Enzyme | Total peptides | Mean length | Median length | 7-35aa range peptides | Sequence coverage (%) | Isoform-unique peptides |
| --- | --- | --- | --- | --- | --- | --- |
| Trypsin/Chymotrypsin | 4,053,121 | 10.3 | 7 | 1,907,193 | 88.7 | 545,769 |
| Trypsin/Asp-N | 3,928,784 | 10.6 | 7 | 1,859,247 | 88.2 | 525,351 |
| Trypsin/Lys-C | 3,017,079 | 13.8 | 7 | 1,442,574 | 78.6 | 408,117 |
| Trypsin-Lys-C | 2,455,307 | 8.5 | 6 | 1,042,209 | 69.6 | 290,657 |
| Trypsin | 2,377,488 | 8.8 | 6 | 1,028,635 | 69.5 | 287,199 |
| Glu-C | 2,599,479 | 8.0 | 5 | 1,049,420 | 67.8 | 287,602 |
| Chymotrypsin | 1,675,637 | 12.4 | 8 | 878,560 | 64.1 | 258,574 |
| Asp-N | 1,562,709 | 13.3 | 9 | 834,695 | 63.3 | 239,505 |
| Trypsin-Asp-N | 3,735,120 | 5.6 | 4 | 1,076,459 | 59.7 | 292,529 |
| Trypsin-Chymotrypsin | 4,015,741 | 5.2 | 4 | 1,078,818 | 59.2 | 291,116 |
| Lys-N | 1,380,762 | 15.1 | 9 | 719,633 | 56.0 | 207,805 |
| Lys-C | 1,377,666 | 15.1 | 9 | 719,763 | 56.0 | 207,580 |
| Arg-C | 1,173,294 | 17.8 | 11 | 620,534 | 50.4 | 182,158 |

It is not surprising that the total numbers of digested peptides are higher for protease combinations than they are for single proteases, due to combining multiple sets of peptides in the case of parallel digests (‘/’) or increased cleavage sites by using multiple proteases in concurrent digests (‘-’). Despite generating a high number of unfiltered peptides, the concurrent Trypsin-Asp-N and Trypsin-Chymotrypsin combinations give relatively low sequence coverage % in comparison to the other digests, due to many digested peptides being shorter than 7 amino acids and therefore below the filtering threshold. The concurrent Trypsin-Lys-C combination gives a slight increase in sequence coverage (0.4%) when compared to the single tryptic digest, likely due to Lys-C cleaving after Proline, whereas trypsin alone does not (Keil, 1992), and therefore increasing cleavage frequency. This combination is favoured *in vitro* because while both proteases cleave at lysine, Lys-C is more efficient at lysine cleavage than trypsin, and therefore combining them reduces the number of mis-cleaved peptides (Glatter et al., 2012). It should be noted that the order in which the enzymes are added to a sample concurrently doesn’t affect the results *in silico*, however this may not be the case *in vitro*.

#### ii. Selecting enzymes with the highest coverage of specific residues

Amino acid composition differs across protein types and families, and the level of post-translational modification differs between amino acids. For example, lysine and arginine, the residues that trypsin cleaves at, are enriched 2.37-fold and 1.95-fold at exon-exon junctions (Wang et al., 2018), but are less frequent in membrane proteins (Kyte and Doolittle, 1982). Acetylation is a PTM that typically occurs on lysine residues. Being able to plan a digest around maximising the coverage of a specific residue may prove useful and options to filter for peptides containing a specific amino acid and calculating amino acid coverage are included in ProtView, which is shown as the percentage of an amino acid in the original sequence that is covered by peptides after filtering for length.

Table 6 shows residue coverage of Cysteine (C), Serine (S), and Lysine (K) from *in-silico A. thaliana* digests to exemplify how much residue coverage % can differ between digest schemes. The digest scheme that gives the highest lysine coverage is Trypsin/Asp-N (80.6%), followed by Trypsin/Chymotrypsin (78.6%), with GluC giving the highest lysine coverage out of the single protease digests (72.8%). This suggests that any of these digests may be favourable to use in analyses focused on the study of acetylation, one of the PTMs that is associated with Lysine. Acetylated Lysine sites are typically not cleaved by trypsin (Garcia et al., 2007), thus peptide length distributions also need to be examined if considering Trypsin/Asp-N or Trypsin/Chymotrypsin. For all three of the residues examined here, parallel protease combinations give the highest coverages, with the exception of Trypsin/Lys-C giving lower lysine coverage (59.1%) due to both of these proteases cleaving at lysine. It should be noted that the protease that gives the highest residue coverage *in silico* may be otherwise unsuitable for use in a certain analysis or require adaptations to the experimental design *in vitro*. For example, in the context of ubiquitination, and despite Glu-C giving high lysine coverage, Glu-C digestion will result in peptides with a STLHLVLRLRGG ubiquitin remnant attached to Lys, causing a +1302.79 Da mass shift that needs to be taken into account in the database search (Warren et al., 2005).

**Table 6.** Residue coverage statistics for A. thaliana using Cysteine (C), Serine (S), and Lysine (K)

| enzyme | total peptides | mean length | filtered peptides (7-35 aa) | protein sequence coverage (%) | C coverage | S coverage | K coverage |
| --- | --- | --- | --- | --- | --- | --- | --- |
| Arg-C | 1,173,294 | 17.8 | 620,534 | 50.4 | 51.5 | 50.1 | 51.1 |
| Asp-N | 1,562,709 | 13.3 | 834,695 | 63.3 | 51.7 | 63.1 | 65.9 |
| Chymotrypsin | 1,675,637 | 12.4 | 878,560 | 64.1 | 68.6 | 63.4 | 63.4 |
| Glu-C | 2,599,479 | 8.0 | 1,049,420 | 67.8 | 74.6 | 71.7 | 72.8 |
| Lys-C | 1,377,666 | 15.1 | 719,763 | 56.0 | 58.1 | 54.9 | 52.3 |
| Lys-N | 1,380,762 | 15.1 | 719,633 | 56.0 | 58.1 | 59.9 | 52.2 |
| Trypsin | 2,377,488 | 8.8 | 1,028,635 | 69.5 | 75.1 | 72.7 | 44.6 |
| Trypsin/Asp-N | 3,928,784 | 10.6 | 1,844,116 | 87.5 | 87.7 | 88.5 | 80.6 |
| Trypsin/Chymotrypsin | 4,053,121 | 10.3 | 1,889,594 | 88.0 | 91.9 | 88.9 | 78.6 |
| Trypsin/Lys-C | 3,017,079 | 13.8 | 1,432,830 | 77.9 | 82.1 | 79.0 | 59.1 |
| Trypsin-Asp-N | 3,735,120 | 5.6 | 1,076,459 | 59.7 | 31.9 | 67.8 | 30.7 |
| Trypsin-Chymotrypsin | 4,015,741 | 5.2 | 1,078,818 | 59.2 | 66.0 | 67.6 | 28.7 |
| Trypsin-Lys-C | 2,455,307 | 8.5 | 1,042,209 | 69.6 | 75.3 | 73.2 | 42.3 |

### Using ProtView to analyse digest outcomes in a transcriptomic context

ProtView provides unique opportunities to examine transcriptomic regulations using proteomic evidence by mapping the digested peptides to the genome reference. ProtView identifies junction-covering peptides and provides junction summary information for each digest scheme. This information is shown in Table 7, consisting of the number of junction-covering peptides, number of junctions covered, unique junctions covered (to avoid double counting of junctions shared between isoforms), and the percentage of junctions that are covered by peptides from each digest after filtering. The function for counting isoform-unique peptides mentioned previously can be appended to the junction summary table if being examined in a transcriptomic context. The number of isoform-unique peptides was calculated for both the entire sets of peptides generated and the junction-covering peptides.

**Table 7.** Junction Summary statistics generated for A. thaliana using ProtView

| Enzyme | Junction spanning peptides | Unique junctions covered | Total junctions covered | Total junction coverage (%) | Isoform-unique peptides | Junction covering isoform-unique peptides |
| --- | --- | --- | --- | --- | --- | --- |
| Trypsin/Chymotrypsin | 286442 | 101973 | 201015 | 84.5 | 545769 | 65589 |
| Trypsin/Asp-N | 276676 | 99323 | 195792 | 82.3 | 525351 | 63561 |
| Trypsin/Lys-C | 207385 | 80087 | 156990 | 66.0 | 408117 | 48464 |
| Chymotrypsin | 152914 | 77761 | 152914 | 64.2 | 258574 | 34697 |
| Glu-C | 147618 | 74684 | 147618 | 62.0 | 287602 | 33211 |
| Asp-N | 143447 | 72751 | 143447 | 60.3 | 239505 | 32777 |
| Trypsin | 133528 | 68070 | 133528 | 56.1 | 287199 | 30892 |
| Trypsin-Lys-C | 133484 | 68079 | 133484 | 56.1 | 290657 | 30917 |
| Lys-N | 129305 | 65925 | 129305 | 54.3 | 207805 | 29634 |
| Trypsin-Asp-N | 115813 | 58815 | 115813 | 48.7 | 292529 | 26358 |
| Trypsin-Chymotrypsin | 114895 | 58311 | 114895 | 48.3 | 291116 | 26014 |
| Lys-C | 111835 | 57631 | 111835 | 47.0 | 207580 | 26656 |
| Arg-C | 106708 | 54338 | 106708 | 44.8 | 182158 | 24673 |

Table 7 exemplifies the format of a junction summary table generated by ProtView, with the digest schemes sorted in order of highest to lowest junction coverage %. The output shows that in terms of single enzyme digests, Chymotrypsin, Glu-C, and Asp-N outperform trypsin in terms of the number of junction-covering peptides generated and junctions covered for the Arabidopsis proteome, further underlining the point that trypsin may not always be the most optimal choice. If carrying out an analysis where maximising splice junction coverage is a priority, these example results suggest that combining chymotryptic peptides in parallel with tryptic peptides can theoretically give a 28.4% increase in junction coverage compared to trypsin alone, in addition to giving the most isoform-unique peptides that can be used to discriminate between protein isoforms.

In addition to the aforementioned overview statistics, ProtView can map peptides onto the genome, allowing the downstream examination and visualization of digested peptides to identify post-transcriptional regulations. For example, AT1G18390 is a gene with two transcript/protein isoforms with alternative transcription start sites. Visualisation of AT1G18390 *in-silico* generated peptides on the genome (Figure 4) shows coverage of the alternative transcriptional start sites (vertical dashed lines) by peptides and allows for isoform-specific and exon-exon junction covering peptides to easily be identified. In this example, trypsin and Asp-N generate peptides that cover the transcription start sites in both isoforms, while the peptides generated by Lys-C do not cover the transcription start site of the AT1G18390.1 isoform. Similarly, AT5G45830 is a gene on the negative strand with alternative stop sites that visually examined on the genome (Figure 5).

**Figure 4.**
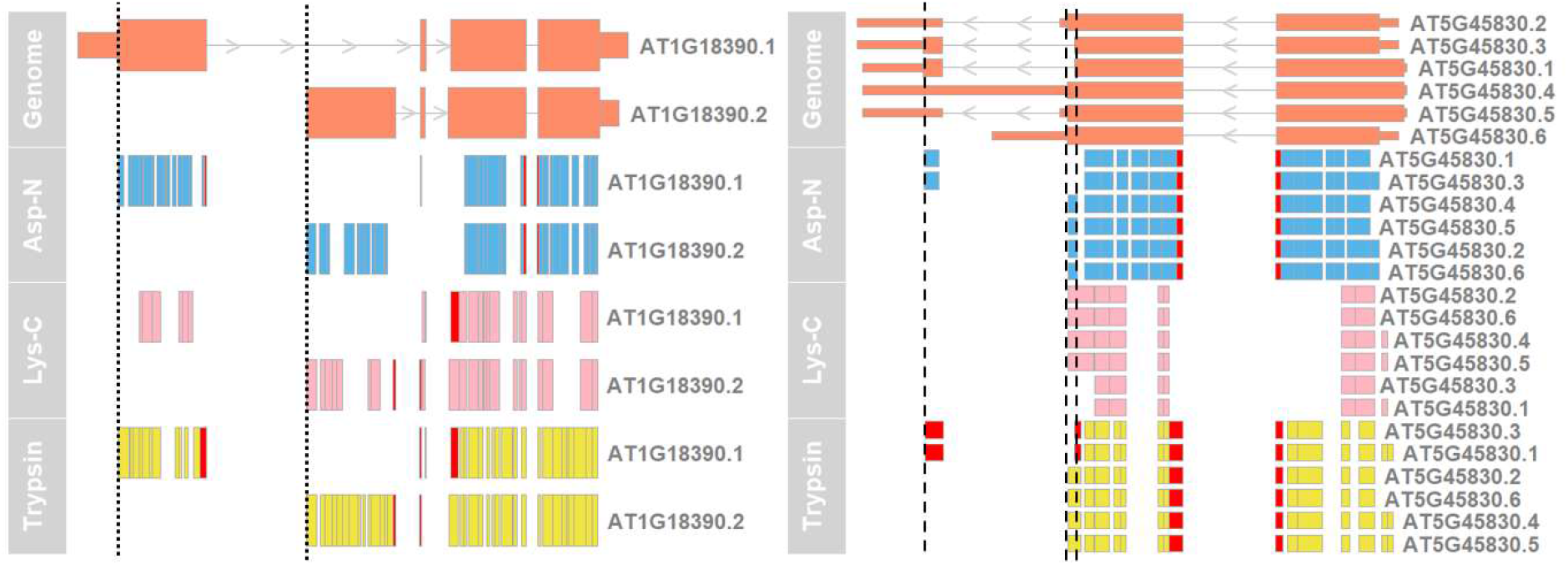
AT1G18390 and AT5G45830 in-silico generated peptides mapped onto the genome. The first row shows the genome; thick coloured boxes represent exons, thin coloured boxes represent UTRs, grey lines linking between exons represent introns, and vertical lines show alternative start(AT1G18390) (dotted) or stop((AT5G45830) (dashed) sites. The subsequent rows represent peptides mapped onto the genome, with exon-exon junction-covering peptides in red.

**Figure 5.**
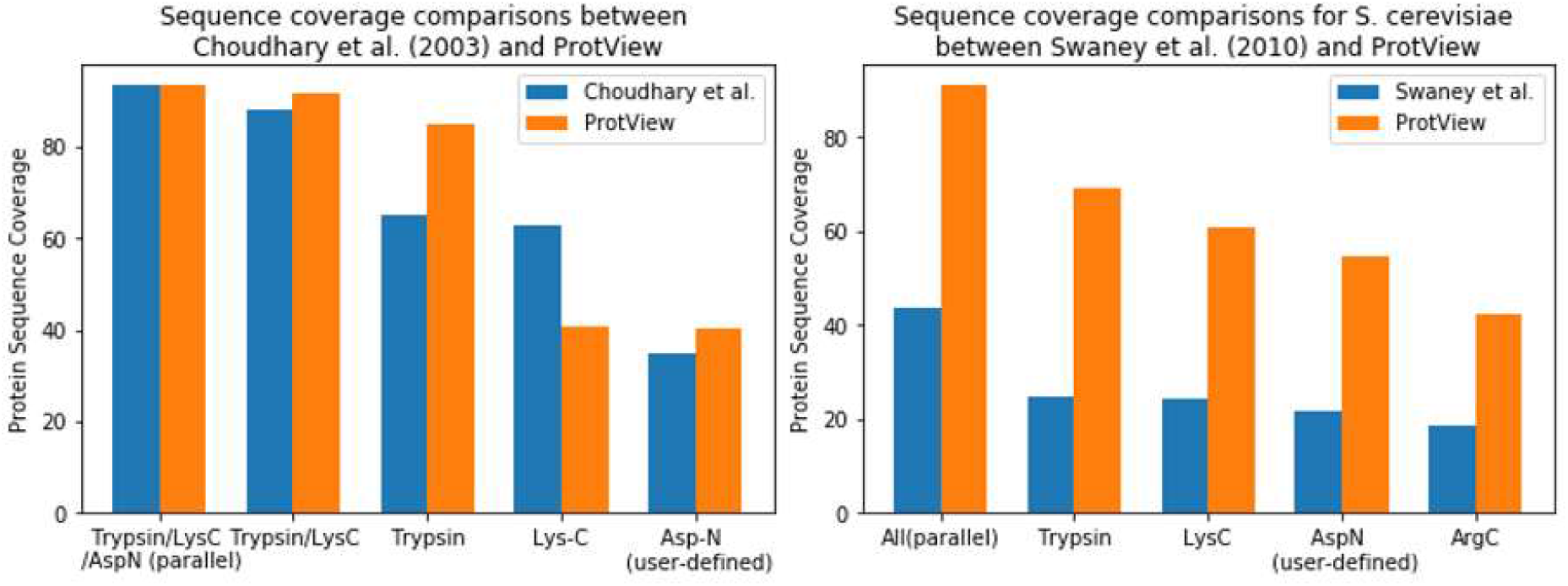
sequence coverage comparisons for human recombinant tissue plasminogen activator between Choudhary et al. (2003) and ProtView and S. cerevisiae between Swaney et al. (2010) and ProtView.

### Comparing ProtView in silico results with in vitro experimental data

To evaluate how representative the order of protein sequence coverage given by ProtView for different digest schemes is of an *in vitro* experiment, comparisons were carried out between ProtView and two publications that compare protein sequence coverage using different proteases. Choudhary et al. (2003) examined a human recombinant tissue plasminogen activator protein using Trypsin, Lys-C, Asp-N and their parallel combinations. Swaney et al. (2010) examined *S. cerevisiae* digests with Trypsin, Arg-C, Asp-N, Lys-C, and all these proteases in parallel. The proteins and digest schemes used in the publications were digested with RPG and processed with ProtView to give protein sequence coverage percentages for each digest scheme. Protein sequence coverage values were then compared between those obtained experimentally in the publications and the *in silico* generated values.

In the first comparison, the sequence coverage obtained by Choudhary et al. with Asp-N was much lower than predicted (34.9% for Choudhary et al., 67.1% for Protview), due to the RPG cleavage rules for Asp-N erroneously including cleavage at all Cysteines, despite Asp-N actually cleaving at Cysteic acid, a vanishingly rare oxidation form of cysteine in natural protein samples (Paulech et al., 2015). It is highly unlikely therefore that Asp-N would deliver a meaningful number of peptides showing cleavage at “Cysteine”. The regions covered by Choudhary et al. do not show cleavage at Cys, thus a user-defined Asp-N (denoted as Asp-N[~C]) cleaving only at Aspartic Acid, was used to repeat the comparison. The highest sequence coverage is obtained by using parallel enzyme digests, with Trypsin/Lys-C/Asp-N[~C] (93.3% for Choudhary et al., 93.4% for Protview), followed by Trypsin/Lys-C (88.2% and 91.8%), Trypsin (65% and 85%), Lys-C (62.8% and 40.6%), and Asp-N[~C] (34.9% and 40.2%). The rankings between the ProtView results and the experimental results are the same, with a spearman correlation of 1 (Figure 5). Two possible explanations were found for the experimental Lys-C coverage being higher than the predicted ProtView value: 1) Peptides above the 35aa filter cutoff length used by ProtView were identified experimentally due to being within the mass range used in the database search. 2) Non-specific cleavage of Lys-C, which can occasionally occur *in vitro* despite Lys-C being highly specific (Raijmakers et al., 2010), but is not considered in the *in-silico* predictions..

In the comparison to Swaney et al. (2010), using user-defined Asp-N rules (Asp-N[~C]), all enzymes in parallel achieved the highest sequence coverage (43.4% for Swaney et al., 91.0% for ProtView), followed by Trypsin (24.5%, 68.9%), Lys-C (24.3%, 60.8%), Asp-N[~C] (21.5%, 54.7%) and ending with Arg-C providing the lowest coverage (18.6%, 42.2%). As in the previous comparison, the spearman correlation value is 1.. The comparisons shows that ProtView can correctly predict the ranking of protease performance and provide rapid pre-analysis to assist in the choice of proteases for addressing a given experimental question.

## CONCLUSIONS

Evaluation of digest schemes *in silico* can save on time and resources compared to *in vitro* evaluations. ProtView is a novel software tool for the evaluation of digest schemes, designed to process and analyse *in-silico* digest output by different enzymes and their combinations in multiple contexts. It is clear from our analysis that depending on the focus of the investigation, the ideal choice of enzyme could vary considerably. The enzyme combinations that provide the best protein sequence coverage do not necessarily provide the best view of the proteome in terms of specific PTMs; Therefore, ProtView is timely to provide a tailored analysis that facilitates the decision and experimental planning process. Preliminary validations show that ProtView can reliably predict the majority of experimentally determined protein sequence coverage orders between digest schemes. ProtView is the first tool that maps the peptides to the reference genome, which allows transcriptomic activities, such as transcriptional (alternative transcriptional start sites and stop sites) and post-transcriptional regulations (alternative splicing), to be studied using proteomic experimental evidence. Mapping peptides to the genome also creates the possibility in the future to integrate sequence variations of different species and their sub-species (e.g cultivars/eco-types/landraces in plants), to derive a list of individualized peptides that is possible to be detected in mass spectrometry-based proteomics experiments.

## Supplementary Material

**Supplementary Figure 1.**
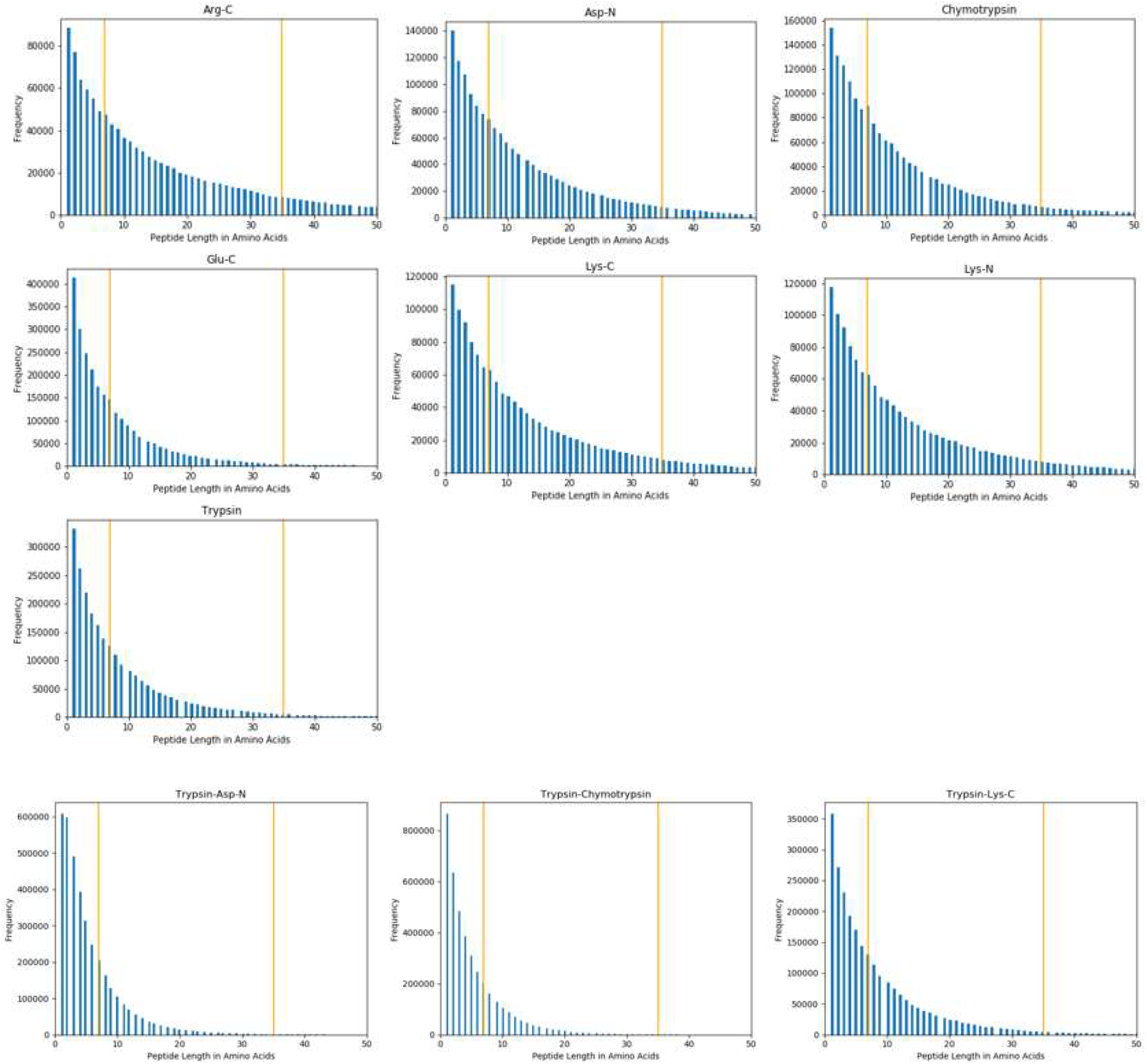
Length distributions of A. thaliana peptides generated by single enzyme digests, showing lengths up to 50 amino acids. Orange vertical lines indicate the 7-35 amino acid length range used in these analyses and shown in Table 6.

**Supplementary Table 1.** Cleavage specificities of the enzymes used in RPG digests in the benchmark analyses in this article

| Residue | R |  | C |  | D |  | F |  | Y |  | W |  | E |  | K |  | Exceptions |
| --- | --- | --- | --- | --- | --- | --- | --- | --- | --- | --- | --- | --- | --- | --- | --- | --- | --- |
| Enzyme \ Terminal | C | N | C | N | C | N | C | N | C | N | C | N | C | N | C | N |  |
| ArgC | X |  |  |  |  |  |  |  |  |  |  |  |  |  |  |  |  |
| Asp-N |  |  |  | X |  | X |  |  |  |  |  |  |  |  |  |  |  |
| Chymotrypsin<br>(high specificity) |  |  |  |  |  |  | X |  | X |  | X |  |  |  |  |  | FP, YP, WP,<br>WM |
| GluC |  |  |  |  | X |  |  |  |  |  |  |  | X |  |  |  |  |
| Lys-C |  |  |  |  |  |  |  |  |  |  |  |  |  |  | X |  |  |
| LysN |  |  |  |  |  |  |  |  |  |  |  |  |  |  |  | X |  |
| Trypsin | X |  |  |  |  |  |  |  |  |  |  |  |  |  | X |  | KP, RP,<br>CKD, DKD,<br>CKH, CKY,<br>CRK, RRH,<br>RRR |

**Supplementary Table 2.** Summary statistics of in silico digests carried out on A. thaliana for all proteases and pairwise protease combinations (‘/’ for parallel, ‘-’ for concurrent)

| Enzyme | Total peptides | Mean length | Median length | Filtered peptides (7-35 aa) | Protein sequence coverage (%) | Isoform unique peptides |
| --- | --- | --- | --- | --- | --- | --- |
| Arg-C | 1173294 | 17.8 | 11 | 620534 | 50.4 | 182158 |
| Arg-C/Asp-N | 2730374 | 15.3 | 10 | 1452634 | 81.4 | 420689 |
| Arg-C/Chymotrypsin-high | 2848912 | 14.6 | 9 | 1499091 | 81.6 | 440727 |
| Arg-C/Glu-C | 3772760 | 11.1 | 6 | 1669951 | 83.4 | 469753 |
| Arg-C/Lys-C | 2550939 | 16.4 | 10 | 1340291 | 77.8 | 389723 |
| Arg-C/Lys-N | 2547448 | 16.4 | 10 | 1337483 | 77.8 | 388979 |
| Arg-C/Trypsin | 3009921 | 13.9 | 7 | 1414582 | 78.7 | 401303 |
| Arg-C-Asp-N | 2606920 | 8.0 | 6 | 1108077 | 70.1 | 308649 |
| Arg-C-Chymotrypsin-high | 2800617 | 7.5 | 5 | 1104433 | 69.3 | 308969 |
| Arg-C-Glu-C | 3724458 | 5.6 | 4 | 1064510 | 60.6 | 291623 |
| Arg-C-Lys-C | 2502642 | 8.3 | 5 | 1050942 | 69.7 | 292816 |
| Arg-C-Lys-N | 2424059 | 8.6 | 6 | 1050942 | 69.7 | 294907 |
| Arg-C-Trypsin | 2435752 | 8.6 | 6 | 1054844 | 69.9 | 289667 |
| Asp-N | 1562709 | 13.3 | 9 | 834695 | 63.3 | 239505 |
| Asp-N/Chymotrypsin-high | 3228399 | 12.9 | 8 | 1709104 | 86.4 | 496638 |
| Asp-N/Glu-C | 4135748 | 10.1 | 6 | 1882591 | 83.3 | 526570 |
| Asp-N/Lys-C | 2934267 | 14.2 | 9 | 1551948 | 83.1 | 446174 |
| Asp-N/Lys-N | 2943460 | 14.2 | 9 | 1554321 | 83.1 | 447300 |
| Asp-N-Chymotrypsin-high | 3057552 | 6.8 | 5 | 1026380 | 63.2 | 308789 |
| Asp-N-Glu-C | 3903157 | 5.3 | 3 | 1038222 | 69.6 | 284585 |
| Asp-N-Lys-C | 2799817 | 7.4 | 5 | 1122250 | 67.4 | 304717 |
| Asp-N-Lys-N | 2895153 | 7.2 | 5 | 1094902 | 68.0 | 303757 |
| Chymotrypsin-high | 1675637 | 12.4 | 8 | 878560 | 64.1 | 258574 |
| Chymotrypsin-high/Lys-C | 3053296 | 13.7 | 9 | 1598319 | 84.3 | 466150 |
| Chymotrypsin-high/Lys-N | 3050069 | 13.7 | 9 | 1595540 | 84.3 | 465440 |
| Chymotrypsin-high-Lys-C | 3004985 | 6.9 | 5 | 1120839 | 68.3 | 309886 |
| Chymotrypsin-high-Lys-N | 2909465 | 7.2 | 5 | 1128967 | 68.8 | 311870 |
| Glu-C | 2599479 | 8.0 | 5 | 1049420 | 67.8 | 287602 |
| Glu-C/Chymotrypsin-high | 4275107 | 9.8 | 6 | 1927978 | 87.9 | 546174 |
| Glu-C/Lys-C | 3977135 | 10.5 | 6 | 1769177 | 85.2 | 495169 |
| Glu-C/Lys-N | 396799<br>3 | 10.5 | 6 | 1765807 | 85.2 | 494340 |
| Glu-C/Trypsin | 497696<br>4 | 8.4 | 5 | 2078054 | 90.1 | 574792 |
| Glu-C-Chymotrypsin-high | 422680<br>1 | 4.9 | 3 | 1057850 | 56.7 | 286868 |
| Glu-C-Lys-C | 392882<br>7 | 5.3 | 3 | 1017603 | 58.0 | 277973 |
| Glu-C-Lys-N | 376001<br>2 | 5.5 | 4 | 1026685 | 58.5 | 280424 |
| Glu-C-Trypsin | 494353<br>7 | 4.2 | 3 | 911196 | 48.0 | 247326 |
| Lys-C | 137766<br>6 | 15.1 | 9 | 719763 | 56.0 | 207580 |
| Lys-C/Lys-N | 274457<br>4 | 15.2 | 10 | 1439365 | 57.7 | 415356 |
| Lys-C-Lys-N | 259856<br>8 | 8.0 | 1 | 665792 | 52.1 | 194045 |
| Lys-N | 138076<br>2 | 15.1 | 9 | 719633 | 56.0 | 207805 |
| Lys-N/Trypsin | 373589<br>3 | 11.2 | 7 | 1747316 | 79.5 | 494614 |
| Lys-N-Trypsin | 353495<br>1 | 5.9 | 2 | 976235 | 65.4 | 274177 |
| Trypsin | 237748<br>8 | 8.8 | 6 | 1028635 | 69.5 | 287199 |
| Trypsin/Asp-N | 392878<br>4 | 10.6 | 7 | 1844116 | 87.5 | 525351 |
| Trypsin/Chymotrypsin-high | 405312<br>1 | 10.3 | 7 | 1889594 | 88.0 | 545769 |
| Trypsin/Lys-C | 301707<br>9 | 13.8 | 7 | 1432830 | 78.0 | 408117 |
| Trypsin-Asp-N | 373512<br>0 | 5.6 | 4 | 1076459 | 59.7 | 292529 |
| Trypsin-Chymotrypsin-high | 401574<br>1 | 5.2 | 4 | 1078818 | 59.2 | 291116 |
| Trypsin-Lys-C | 245530<br>7 | 8.2 | 6 | 1042209 | 69.6 | 290657 |

**Supplementary Table 3.** Junction Summary statistics generated for A. thaliana using ProtView for all proteases and pairwise protease combinations (‘/’ for parallel, ‘-’ for concurrent)

| enzyme | Junction<br>spanning<br>peptides | Unique<br>junctions<br>covered | Total<br>junctions<br>covered | Total<br>junction<br>coverage<br>(%) | isoform<br>unique<br>peptides |
| --- | --- | --- | --- | --- | --- |
| Arg-C | 106708 | 54338 | 106708 | 44.8 | 182158 |
| Arg-C/Asp-N | 249948 | 93812 | 184954 | 77.7 | 420689 |
| Arg-C/Chymotrypsin-high | 259622 | 96985 | 191119 | 80.3 | 440727 |
| Arg-C/Glu-C | 254326 | 95721 | 189126 | 79.5 | 469753 |
| Arg-C/Lys-C | 218543 | 86269 | 169321 | 71.1 | 389723 |
| Arg-C/Lys-N | 235754 | 90064 | 177340 | 74.5 | 388979 |
| Arg-C/Trypsin | 211060 | 83177 | 163743 | 68.8 | 401303 |
| Arg-C-Asp-N | 154222 | 77981 | 154222 | 64.8 | 308649 |
| Arg-C-Chymotrypsin-high | 154947 | 78335 | 154947 | 65.1 | 308969 |
| Arg-C-Glu-C | 124541 | 63249 | 124541 | 52.3 | 291623 |
| Arg-C-Lys-C | 133636 | 68140 | 133636 | 56.2 | 292816 |
| Arg-C-Lys-N | 149667 | 75662 | 149667 | 62.9 | 294907 |
| Arg-C-Trypsin | 133794 | 68187 | 133794 | 56.2 | 289667 |
| Asp-N | 143447 | 72751 | 143447 | 60.3 | 239505 |
| Asp-N/Chymotrypsin-high | 295886 | 103107 | 203497 | 85.5 | 496638 |
| Asp-N/Glu-C | 290866 | 97271 | 192445 | 80.9 | 526570 |
| Asp-N/Lys-C | 255045 | 94945 | 186872 | 78.5 | 446174 |
| Asp-N/Lys-N | 272752 | 98233 | 193629 | 81.4 | 447300 |
| Asp-N-Chymotrypsin-high | 154709 | 77909 | 154709 | 65.0 | 308789 |
| Asp-N-Glu-C | 136795 | 69302 | 136795 | 57.5 | 284585 |
| Asp-N-Lys-C | 139246 | 70694 | 139246 | 58.5 | 304717 |
| Asp-N-Lys-N | 150661 | 76080 | 150661 | 63.3 | 303757 |
| Chymotrypsin-high | 152914 | 77761 | 152914 | 64.3 | 258574 |
| Chymotrypsin-high/Lys-C | 264749 | 98326 | 193618 | 81.4 | 466150 |
| Chymotrypsin-high/Lys-N | 281926 | 101082 | 199347 | 83.8 | 465440 |
| Chymotrypsin-high-Lys-C | 140548 | 71165 | 140548 | 59.1 | 309886 |
| Chymotrypsin-high-Lys-N | 153581 | 77369 | 153581 | 64.4 | 311870 |
| Glu-C | 147618 | 74684 | 147618 | 62.0 | 287602 |
| Glu-C/Chymotrypsin-high | 300532 | 104325 | 205863 | 86.5 | 546174 |
| Glu-C/Lys-C | 259453 | 97618 | 192543 | 80.9 | 495169 |
| Glu-C/Lys-N | 276607 | 99195 | 195916 | 82.3 | 494340 |
| Glu-C/Trypsin | 281146 | 103259 | 204067 | 85.7 | 574792 |
| Glu-C-Chymotrypsin-high | 120575 | 61040 | 120575 | 50.7 | 286868 |
| Glu-C-Lys-C | 109317 | 55888 | 109317 | 45.9 | 277973 |
| Glu-C-Lys-N | 119523 | 60743 | 119523 | 50.2 | 280424 |
| Glu-C-Trypsin | 84342 | 43214 | 84342 | 35.4 | 247326 |
| Lys-C | 111835 | 57631 | 111835 | 47.0 | 207580 |
| Lys-C/Lys-N | 241140 | 68533 | 134210 | 56.4 | 415356 |
| Lys-C-Lys-N | 106737 | 55000 | 106737 | 44.8 | 194045 |
| Lys-N | 129305 | 65925 | 129305 | 54.3 | 207805 |
| Lys-N/Trypsin | 262789 | 89875 | 177200 | 74.5 | 494614 |
| Lys-N-Trypsin | 127429 | 64907 | 127429 | 53.5 | 274177 |
| Trypsin | 133528 | 68070 | 133528 | 56.1 | 287199 |
| Trypsin/Asp-N | 276676 | 99323 | 195792 | 82.3 | 525351 |
| Trypsin/Chymotrypsin-high | 286442 | 101973 | 201015 | 84.5 | 545769 |
| Trypsin/Lys-C | 207385 | 80087 | 156990 | 66.0 | 408117 |
| Trypsin-Asp-N | 115813 | 58815 | 115813 | 48.7 | 292529 |
| Trypsin-Chymotrypsin-high | 114895 | 58311 | 114895 | 48.3 | 291116 |
| Trypsin-Lys-C | 133484 | 68079 | 133484 | 56.1 | 290657 |

## AUTHOR INFORMATION

Corresponding Author

Present Address

Author Contributions - CRediT Statement

Sophia Puliasis: Methodology, Software, Writing – original draft, Formal analysis, Validation, Visualisation, Conceptualization. Runxuan Zhang: Conceptualization, Methodology, Supervision, Writing – original draft /Writing – review & editing, Project administration, Validation, Resources, Funding Acquisition. Piers Hemsley: Conceptualization, Supervision, Writing – review & editing, Resources, Funding Acquisition. Dominika Lewandowska: Conceptualization, Supervision, Writing – review & editing.

